# The SAMBA tool uses long reads to improve the contiguity of genome assemblies

**DOI:** 10.1101/2021.10.21.465348

**Authors:** Aleksey V. Zimin, Steven L. Salzberg

## Abstract

Third-generation sequencing technologies can generate very long reads with relatively high error rates. The lengths of the reads, which sometimes exceed one million bases, make them invaluable for resolving complex repeats that cannot be assembled using shorter reads. Many high-quality genome assemblies have already been produced, curated, and annotated using the previous generation of sequencing data, and full re-assembly of these genomes with long reads is not always practical or cost-effective. One strategy to upgrade existing assemblies is to generate additional coverage using long-read data, and add that to the previously assembled contigs. SAMBA is a tool that is designed to scaffold and gap-fill existing genome assemblies with additional long-read data, resulting in substantially greater contiguity. SAMBA is the only tool of its kind that also computes and fills in the sequence for all spanned gaps in the scaffolds, yielding much longer contigs. Here we compare SAMBA to several similar tools capable of re-scaffolding assemblies using long-read data, and we show that SAMBA yields better contiguity and introduces fewer errors than competing methods. SAMBA is open-source software that is distributed at https://github.com/alekseyzimin/masurca.

## Introduction

The development of long-read sequencing technologies has revolutionized the genome assembly landscape. It is possible now, for example, to get reads that are up to a million bases long from Oxford Nanopore’s MinION and PromethION instruments (Nurk et al., 2021). These “ultralong” reads are extremely helpful for assembling genomic regions filled with long, complex repeats. However, the high error rates are still an obstacle to obtaining high-quality genome sequences from long reads alone. One effective strategy for creating assemblies with long reads is to generate both short and long reads, and then to assemble the “easy” parts of the genome with short reads and use the long reads to resolve the hard-to-assemble regions. However, for many genomes that have already been sequenced, existing assemblies already capture most of the non-repetitive sequence, obviating the need to generate additional short-read data. For these genome assemblies, adding long read data might provide a fast, cost-effective way to improve their contiguity.

A hybrid approach for scaffolding genomes using long-read data was first proposed in Bashir et al. (2012). The underlying algorithm is straightforward: first it maps long reads to contigs, then it uses the alignments to link the contigs into a graph, and finally it traverses the resulting graph to produce scaffolds, in which gap sizes are estimated from the linking information. In practice, implementing this approach may be quite complex, due to repetitive alignments that introduce loops and nested loops in the graph, and to erroneous links that result from spurious alignments of reads.

In this paper we describe a novel tool called SAMBA (Scaffolding Assemblies with Multiple Big Alignments) that uses long reads to re-scaffold contigs from an existing genome assembly and to fill gaps in those scaffolds. The tool is designed to use 10-30x coverage in long read data with a set of existing contigs, although it can function with lower or higher coverage as well. Several previously developed tools also allow for scaffolding with long reads. These include AHA, which is part of the SMRT software analysis suite (Bashir et al., 2012), SSPACE-LongRead (Boetzer and Pirovano, 2014), and LINKS (Warren et al., 2015), all of which were designed for bacteria and other small genomes. The SMIS scaffolder (https://www.sanger.ac.uk/tool/smis/) utilizes an approach of creating artificial “mate-pairs” from the long reads and using that mate-pair information to scaffold the contigs. The recently published LRScaf tool (Qin et al., 2019) is one of the very few tools besides SAMBA that can handle mammalian-sized genome data, but its algorithm is prone to creating artificial duplications of contigs. All tools except for SMIS attempt to merge contigs when negative gaps (overlaps between contigs within a scaffold) are detected, however none of these tools utilize the consensus of the long reads to fill gaps in the scaffolds as SAMBA does.

The SAMBA tool described here is also a general-purpose assembly improvement tool. It chooses automatically between two modes or operation. It takes as input an assembly, which can be a set of contigs or scaffolds, plus either a set of long reads (mode 1) or another set contigs from the same or a closely related genome (mode 2). It then uses the reads or the other contigs to link and merge contigs of the input assembly. SAMBA is fast and memory efficient: it can scaffold and gap-fill a mammalian genome in only a few hours on a modern 24-32 core server. In this paper we present the results of running SAMBA on data sets from *Arabidopsis thaliana* and human, and we compare its accuracy and run times to other comparable published software.

SAMBA is open source software, distributed with the MaSURCA assembler, included in version 4.0.5 and above.

## Results

We compared the performance of SAMBA with three other long-read scaffolding algorithms: LRScaf version 1.1.11 (Qin et al., 2019), LINKS 1.8.7 (Warren et al., 2015), and SMIS v0.1-alpha (https://www.sanger.ac.uk/tool/smis/). Because AHA and SSPACE were designed for bacterial genomes, their memory (RAM) requirements make them impractical to use for large (mammalian-sized) genomes, so we did not include them in these experiments.

For the experimental comparisons, we used data sets from two organisms, *Arabidopsis thaliana* Ler-0 and human. The Illumina and PacBio data for Arabidopsis was described in (Lee et al., 2014), and the high-quality assembly of these data is available from (Berlin et al., 2015). See Data Availability section for more details. The datasets we used for both Arabidopsis and human experiments are listed in Table 1.

**Table 1.** Data used for the scaffolding experiments, including short and long reads from the model plant *Arabidopsis thaliana*, and reads from two different individual humans. The Illumina reads were paired-end reads of a fixed length. N50 read length is computed by summing all read lengths, and determining the value N50 such that 50% of that total length is covered by reads of N50 or longer.

|  |  |  |  |
| --- | --- | --- | --- |
|  | **Number of reads** | **Genome coverage** | **N50 read length** |
| ***A. thaliana*** |  |  |  |
| **Illumina** | 46,129,480 | 115x | 300 |
| **PacBio P6-C4** | 3,448,228 | 118x | 7,205 |
| ***H. sapiens* CHM13** |  |  |  |
| **Oxford Nanopore** | 29,468,868 | 118x | 56,695 |
| ***H. sapiens* HG01243** |  |  |  |
| **Illumina paired-end** | 1,307,137,030 | 59x | 150 |
| **PacBio CLR** | 23,672,554 | 113x | 28,837 |
| **Oxford Nanopore** | 16,239,849 | 58x | 44,052 |

### Arabidopsis thaliana evaluation

We first assembled the genome of *Arabidopsis thaliana* from Illumina paired-end reads, without using PacBio data, using MaSuRCA 4.0.3 (Zimin et al., 2013, Zimin et al., 2017). For clarity we will refer to this assembly as Athal-Illumina. The Athal-Illumina assembly contained a total of 118,048,454 bp of sequence in 4663 contigs, with an NG50 contig size of 172,147 bp; i.e., the size at which >50% of the assembly is contained in contigs of this length or longer. The NG50 size was computed by Quast (Gurevich et al., 2013) using as reference the previously-published assembly produced with the PacBio data (Berlin et al., 2015), which we refer to as Athal-Berlin. NG50 differs from the standard N50 computation in that the genome size used for the computation is that of the reference (Athal-Berlin) rather than that of Athal-Illumina itself. Further evaluation of the Athal-Illumina assembly with Quast reported an NGA50 of 150,736 bp with 139 mis-assemblies. Quast computes NGA50 by splitting the query (Athal-Illumina) genome at mis-assemblies detected by aligning it to the reference (Athal-Berlin), and then re-running the NG50 computation.

We then scaffolded Athal-Illumina using (1) additional PacBio reads and (2) the Athal-Berlin assembly. For the first re-scaffolding, we randomly chose multiple subsets of long PacBio reads, each representing 10-50x coverage, from the total of 118X coverage available (Table 1). We then used these to re-scaffold the Athal-Illumina assembly, and then to evaluate the scaffolder’s performance as function of long-read coverage. We evaluated assembly quality with Quast before and after scaffolding, using the Athal-Berlin assembly as the reference. For both SAMBA and LRScaf we used similar scaffolding parameters, setting the minimum long-read alignment length to 2500 bp and the maximum overhang length to 1000 bp. We experimented with different values of minimum alignment length, and 2500 seemed to provide the best balance between the number of mis-assemblies and the size of output scaffolds for this data set. Decreasing that number led to more aggressive scaffolding (i.e., larger scaffolds) at the expense of a higher mis-assembly rate for both programs. LINKS and SMIS use different approaches to scaffolding and do not offer the user the ability to adjust these parameters, and thus we used the defaults for these programs.

The scaffolding results for Arabidopsis as a function of long-read coverage depth are shown in Figure 1. SAMBA had the smallest number of scaffolding errors (Fig. 1d), and it yielded the longest contigs (Fig. 1a). LRScaf and SMIS created the longest scaffolds (Fig. 1b). LRScaf had the highest number of errors in both contigs and scaffolds (Fig. 1c,d). Improvements in contiguity obtained by all scaffolders saturated at about 30x coverage by long reads. We also noticed that the total amount of sequence in LRScaf’s output varied substantially for different inputs, ranging from 118–127 Mbp for this 120 Mbp genome. We traced that to a tendency for LRScaf to create erroneous duplicate copies of contigs on the ends of some scaffolds. (Note that we reported this as an issue on the LRScaf github site.)

**Figure 1.**
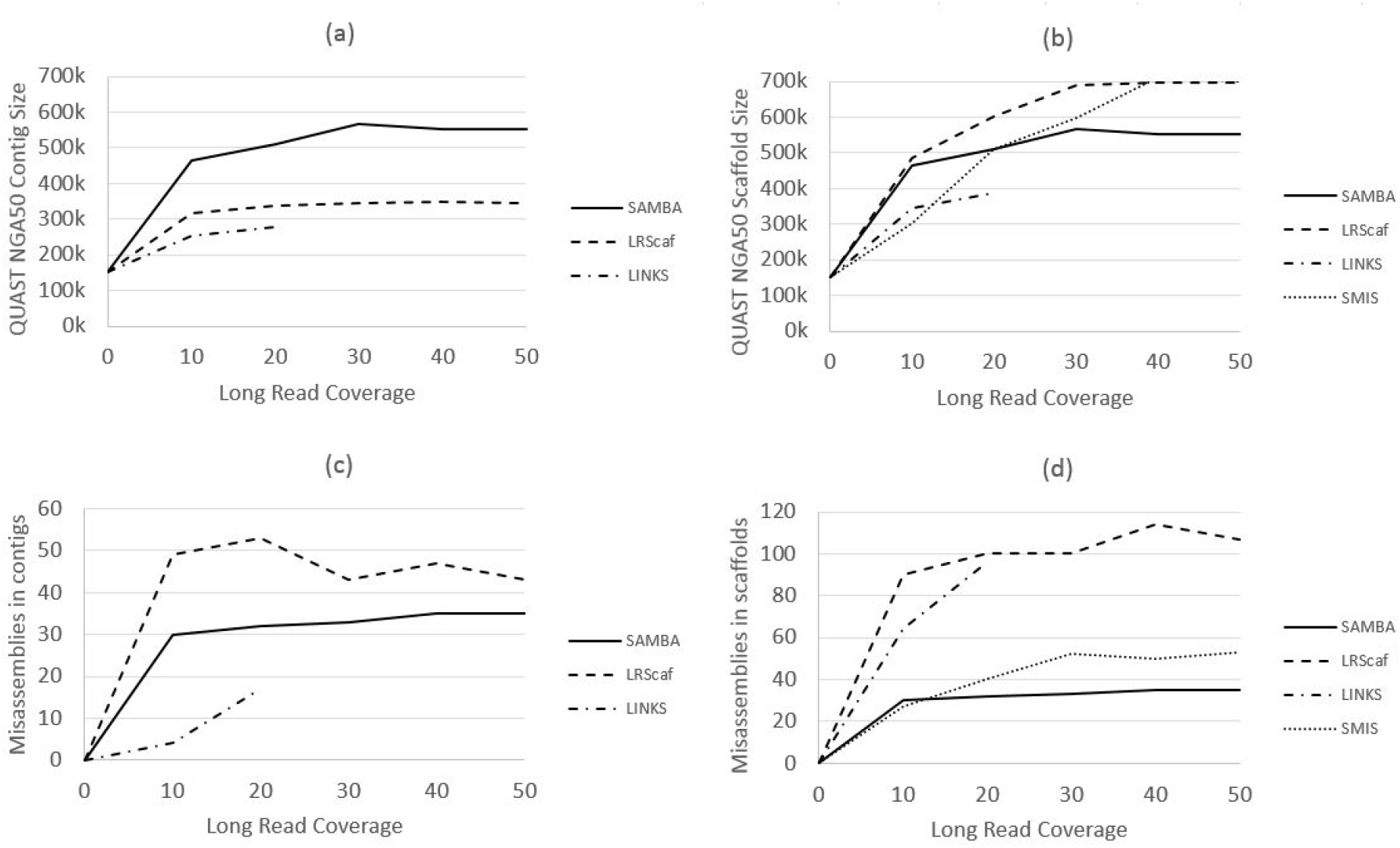
Scaffolding Athal-Illumina assembly with LINKS, SMIS, LRScaf, and SAMBA. We compared performance as a function of long-read (PacBio) coverage, shown on the x-axis. Panels (a) and (b) show the NGA50 size for contigs and scaffolds after scaffolding with each tool. SAMBA outputs contigs, and thus its scaffold and contig results are the same. Panels (c) and (d) show the number of mis-assemblies introduced by the scaffolding process. The LINKS scaffolder required more than 512GB of RAM for PacBio coverage >20x, and thus we did not run it with higher coverage. SMIS does not change contigs, so we did not include its statistics on panels (a) and (c).

When we used the Athal-Berlin assembly (Berlin et al., 2015) to scaffold the Athal-Illumina assembly, we were able to achieve an NGA50 of 1,173 Kbp, more than twice as large as the 567 Kbp that was the best NGA50 length when we re-scaffolded with long reads alone. Not surprisingly, assembly using Illumina and PacBio data was far superior to using Illumina data alone, so it would make more sense to simply use both data sets in the first place. However, if one is assembling a genome that is very closely related to a previously-assembled species, one can use this approach to improve an Illumina-only assembly. SAMBA introduced only one contig mis-assembly, compared to 30 or more mis-assemblies introduced when using the PacBio data with any of the scaffolding programs (Fig. 1d). This shows the effectiveness of SAMBA in reconciling different assemblies of the same genome. Ideally, we would like to achieve the same contiguity as the reference when scaffolding with a better assembly of the same species, but contig mis-assemblies in the Athal-Illumina assembly prevented that here. Below we discuss further the issue of detecting contig mis-assemblies.

### Homo sapiens evaluation

Next, we examined the scaffolders’ performance on human data, which we evaluated on two different human data sets. The first data set is from CHM13, a haploid human cell line that was used to create the first-ever complete human genome assembly (Miga et al., 2020; Nurk et al., 2021), and the second data set is diploid data from a Puerto Rican individual, HG01243, sequenced as part of the human pangenome project (see Data Availability section). Details of these datasets are listed in Table 1. The Telomere-to-Telomere (T2T) consortium has produced a very high-quality, gap-free assembly of CHM13 called CHM13 v1.1 (Nurk et al., 2021) which makes evaluations of assembly correctness for the CHM13 data straightforward. However, due to its pseudo-haploid nature, the CHM13 genome is considerably easier to assemble than a typical diploid human. Thus we also include in our comparison assemblies of the diploid human sample HG01243.

For the first human comparison, we used a draft assembly of CHM13 produced by the Genome Institute at Washington University School of Medicine in 2015 (NCBI accession GCA_000983455.2). This assembly, which we refer to as CHM13-WashU, was produced with the Falcon assembler using 52x coverage in PacBio reads. Evaluation of this assembly with Quast yielded an NGA50 contig size of 9,287,872 bp, with 2,346 apparent mis-assemblies when compared to the CHM13 v1.1 assembly. To re-scaffold the CHM13-Washu assembly, we used the ONT reads from the CHM13hTERT cell line described in Table 1.

For the comparison on diploid human data, we produced an assembly of HG01243 from Illumina and PacBio data (Table 1) using MaSuRCA, which we refer to as HG01243-IP. Evaluation of this assembly with Quast yielded an NG50 contig size of 9,756,965 bp with 2,431 apparent mis-assemblies when compared to CHM13 v1.1. We then re-scaffolded the HG01243-IP assembly with ONT ultralong reads. We note that because the reference here was from a different individual, some of the mis-assemblies detected by Quast are likely to be genuine structural differences between the two individuals, so the number of mis-assemblies is an overestimate, and the NGA50 contig size (NG50 after the contigs are split at the apparent mis-assembly locations) is an underestimate. However, we can still use the number of mis-assemblies detected by Quast for a relative comparison of the different scaffolders.

We then used SAMBA, LINKS, SMIS and LRScaf to scaffold the CHM13-WashU and HG10243-IP assemblies using randomly chosen subsets of ultralong ONT reads representing 10x, 20x and 30x coverage. We set the minimum long read alignment to 5000 bp for both SAMBA and LRScaf. LINKS required more than 512 GB of computer memory even at the lowest level of ONT read coverage, and SMIS was unable to complete the scaffolding in 2 weeks on a 24-core server; therefore, we did not include the results from either of these tools in these comparisons. We evaluated contiguity and correctness with Quast using CHM13 v1.1 as the reference.

Figure 2 compares the resulting scaffold sizes as a function of read coverage for both assemblies, using NGA50 size for CHM13 and NG50 size for HG01243. Panels (a) and (c) refer to HG01243 evaluations and panels (b) and (d) refer to CHM13 evaluations. For the CHM13 genome, LRScaf produced bigger scaffolds with more errors than SAMBA, while for the HG01243 assembly SAMBA outperformed LRScaf in both scaffold size and the number of errors. It is worth noting here that in the SAMBA output, scaffolds have no gaps because SAMBA fills all scaffold gaps with consensus sequence.

**Figure 2.**
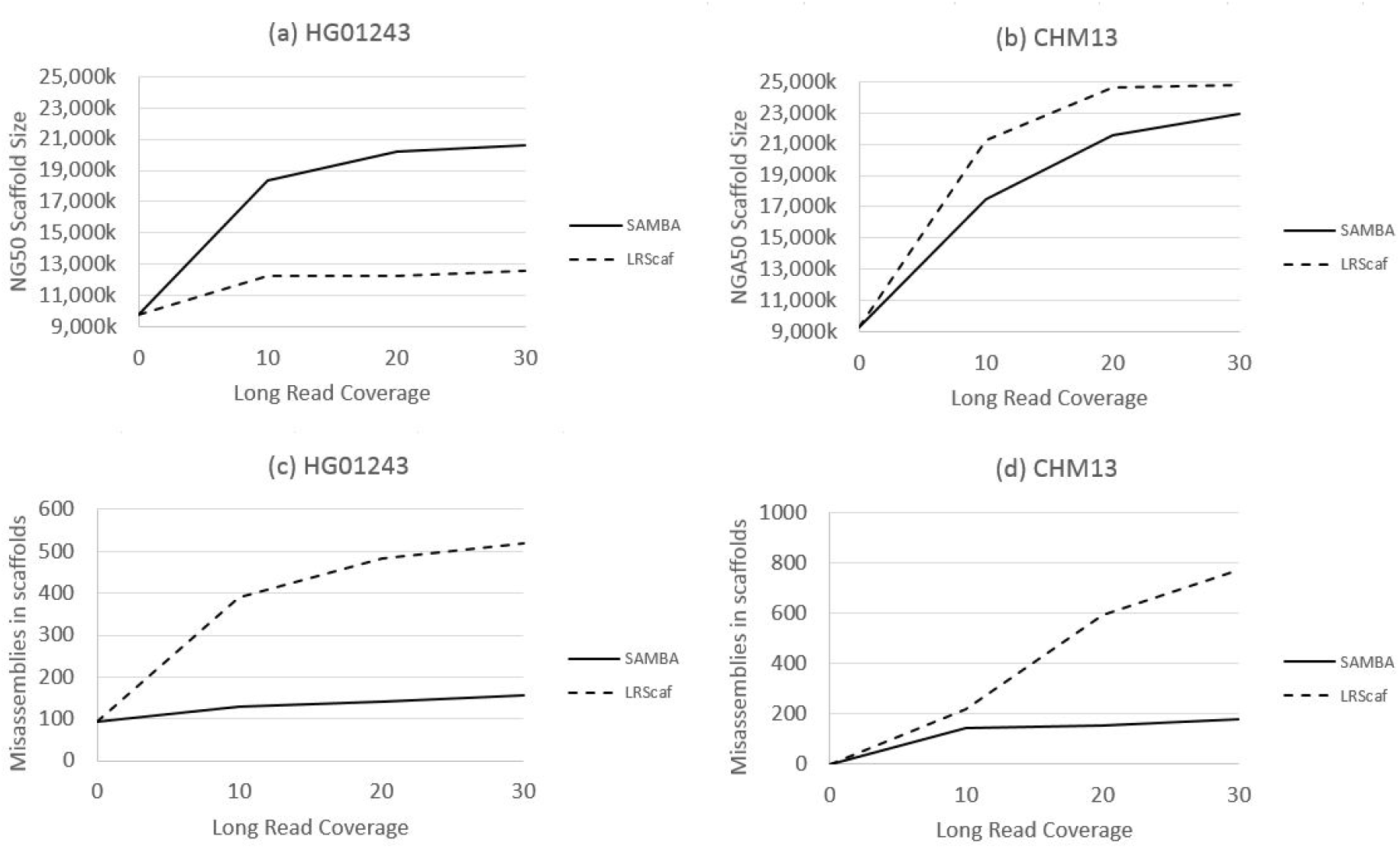
Contiguity and accuracy results for SAMBA and LRScaf on scaffolding of assemblies of human CHM13 (CHM13-WashU) and HG01243 (HG01243-IP) cell lines using varied coverage of additional Oxford Nanopore (ONT) ultralong reads. The x-axis shows coverage in the ultralong ONT reads. All assemblies were evaluated by Quast using CHM13 v1.1 assembly as the reference. Panels (a) and (c) compare NG50 and the number of mis-assemblies for scaffolded HG01243-IP assembly. Panels (b) and (d) show the comparison for CHM13-WashU assembly. LRScaf produces bigger scaffolds for CHM13 data, and SAMBA produces bigger scaffolds for HG01243 data. SAMBA introduces fewer mis-assemblies in all experiments.

In terms of cost-benefit analysis, we observe that scaffold sizes continued to increase as coverage in ONT reads rose from 20x to 30x, suggesting that even deeper coverage would yield some additional improvements.

In all our experiments, SAMBA ran much faster than SMIS and LINKS, but somewhat slower than LRScaf. For example, at 30x coverage in the Arabidopsis experiment (Figure 1), SAMBA scaffolded the Athal-Illumina assembly in about one hour, with 50 minutes spent on computing consensus on the sequence “patches” used for merging contigs. On the same data set, LINKS took nearly 10 hours and SMIS took about 28 hours (Table 2). LRScaf was the fastest, completing the scaffolding in under 5 minutes, with most of the time spent in alignment of the long reads with minimap2. SAMBA does extra work, as compared to LRScaf, in computing the consensus to fill the gaps. When the time spend computing the gap sequences is subtracted, LRScaf and SAMBA have very similar run times. On the human data (Figure 2), SAMBA completed scaffolding and gap filling with 30x Nanopore long reads in 12 hours, with 9 hours spent on initial read alignment by minimap2 and 3 hours spent on computing the consensus sequences to fill gaps. LRScaf finished the scaffolding in about 9 hours. The time spent on scaffolding (resolving the contig graph) was only a few minutes for both LRScaf and SAMBA. For all the timings reported here, we used a 24-core Intel(R) Xeon(R) 6248R server with 512 GB of RAM.

**Table 2.**
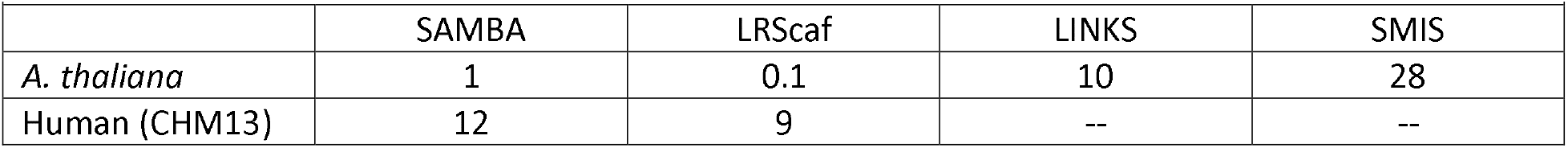
Timings (in hours) that each tool required for scaffolding the *Arabidopsis* and human assemblies with 30x coverage by long reads.

## Methods

The input to SAMBA comprises a set of input contigs C and a set of sequences used to join the contigs together, which we will designate as J. J can contain either long reads or contigs from another assembly of the same organism made, for example, using a different assembler. SAMBA starts off by aligning all sequences J_i_ in J to the contigs C_k_ in C using minimap2 (Li, 2018). We only use the best alignments that are longer than 5Kb (a parameter) that may overhang the end of the contig by no more than 1Kb (also a parameter), and we call these the “proper” alignments. An overhang occurs when an alignment of sequence C_k_ to a sequence J_i_ is such that the alignment stops before an end of C_k_, but J_i_ continues beyond the end of C_k_. The reason why we allow for overhangs is that contigs may have local mis-assemblies on the ends, which frequently are the reason why an assembler could not extend the contig. The requirement of a minimum alignment length ensures that short repeat-induced overlaps are filtered out. We only keep the proper alignments of sequences in J that align to two or more contigs in C, and discard all alignments where J_i_ only aligns properly to a single contig. We note here that in practice SAMBA performs better on relatively contiguous input assemblies, with an N90 >10 Kb; i.e., 90% of the assembled sequence is found in contigs of 10 Kb or longer.

A proper alignment of a sequence J_i_ to two contigs C_i_ and C_j_ in C creates a proper edge between C_i_ and C_j_. If the set J consists of reads that represent relatively deep coverage, we require more than one proper edge between each pair of connected contigs. We then bundle the proper edges and produce consensus sequences for them using the Flye assembler’s consensus module (Kolmogorov et al., 2019). This results in a set of linking sequences. If J represents another assembly of the same or a similar genome (as opposed to a set of long reads), then we skip the consensus step. We then re-align the linking sequences to the contigs in C with minimap2 and build a graph of contigs in C connected by the proper edges, where the edges are the proper alignments of the linking sequences. Each node (each contig) naturally has a beginning and an end with an arbitrary (randomly chosen) direction, and we designate all adjacent edges as either incoming or outgoing.

The next step is to search for linear paths in the graph, defined as sequences of nodes that have exactly one incoming and one outgoing edge. We collapse such paths into super-nodes, as shown in Figure 3. Following this step, we search for “bubbles,” which are paths that branch into two alternative paths and then re-merge. We call the nodes in the bubbles B_1_…B_n_, where n>=2, and all incoming edges to B_1_…B_n_ are outgoing from the same node C_i_ and all outgoing edges are incoming to the same node C_j_. For example, Fig. 3 shows a bubble beginning at node C_2_, which has two outgoing edges, and ending in node C_3_, with two incoming edges. A common source of these bubbles is a haplotype variant in the original diploid genome, which may lead to a contig graph where two haplotypes representing the same location on the genome were assembled into two different contigs. We “pop” the bubbles by choosing the path through the bubble that is the longest of all B_n_ nodes. To resolve “bubbles within bubbles” we perform bubble popping iteratively until no bubbles are left to pop in the graph.

**Figure 3.**
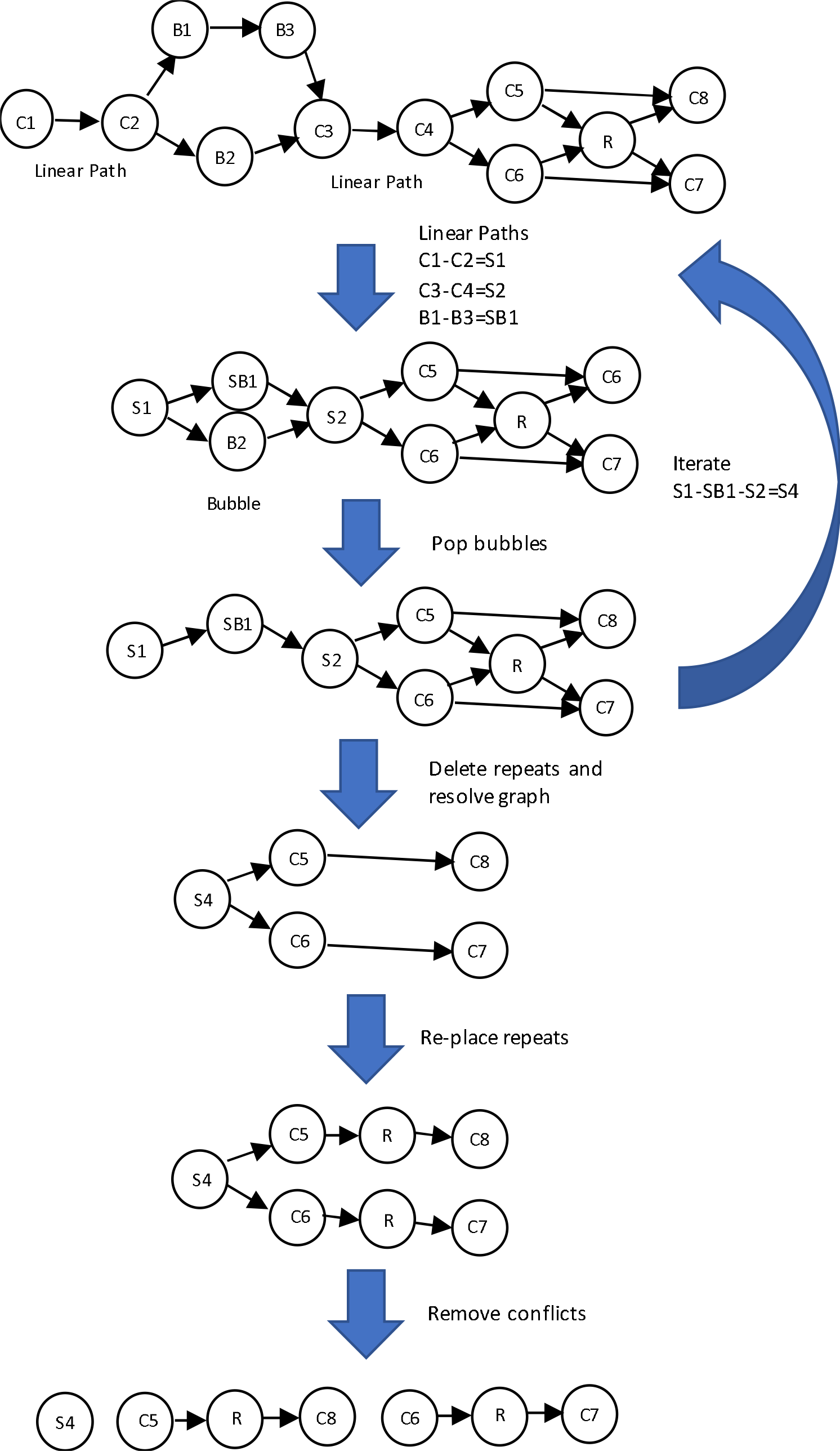
Overview of the SAMBA scaffolding algorithm. The method resolves the initial graph into three final contigs, C1-C2-B1-B3-C3-C4 (S4), C5-R-C8 and C6-R-C7. The repeat contig R is duplicated in the final output. Note that if there were no proper edge between contigs C5 and C8, then the final output would contain C5 and C8 standalone instead of C5-R-C8.

Following bubble popping, we look for repetitive contigs, indicated by nodes or super-nodes that have two or more incoming and two or more outgoing edges. In SAMBA we only aim to resolve repeats that are completely spanned by long reads in J that in turn have long overlaps with unique sequence surrounding the repeat. Therefore, we temporarily remove all repeat nodes from the graph, noting their edges, and re-compute the graph without them. This usually simplifies the graph, introducing breaks in places where there are not enough data to resolve repeats. We then examine all nodes in the resulting graph, and if a node has two or more incoming edges, we delete them both. We do the same if a node has two or more outgoing edges. Thus, for example, if a node has two outgoing edges and a single incoming edge, we keep the single incoming edge and delete the two outgoing edges. This breaks the graph into linear paths. Finally, we re-insert all repeat nodes (contigs) that we temporarily removed into the resulting linear paths according to their edges, duplicating the sequence if necessary. The final linear paths represent the scaffolded contigs. An example of the scaffolding process is depicted in Figure 3. Finally, we collapse the linear paths, and fill positive-sized gaps with the linking sequence obtained from the consensus of linking sequences in J.

Following the scaffolding and gap filling steps, we extract all sequences that SAMBA used to fill gaps in the final scaffolds and align all contigs in C that are shorter than the minimum alignment length threshold (5000bp by default) to these sequences with minimap2. We then delete all short contigs in C that are contained for at least 95% of their length in these gap-filling sequences. This ensures that we do not spuriously duplicate sequences that were originally present in the assembly in the form of short contigs.

## Discussion

We have shown here that SAMBA is a highly effective tool for improving the contiguity of genome assemblies with additional long read data. The primary limitation of SAMBA is determined by the contig lengths in the assembly being scaffolded. Since the default minimum overlap for long reads is 5000 bp, any contigs that are shorter than this minimum will not be scaffolded by SAMBA. We recommend applying SAMBA to assemblies where most of the sequence is in contigs that are longer than 5000 bp, which is not difficult to achieve with modern sequencing technologies and assembly software.

Our future plans for SAMBA development include detection and correction of contig mis-assemblies in the input assembly. We may be able to use discordant alignments, such as cases where long reads map to the interiors of two different contigs, to detect and break mis-assembled contigs. This may allow SAMBA to produce a re-scaffolded genome that is not only more contiguous, but also more accurate than the original assembly.

## Data Availability

The Illumina and PacBio data for *A. thaliana* are available from NCBI SRA under accessions SRX533607, SRX533608 and from http://schatzlab.cshl.edu/data/ectools/. The human CHM13 data is available from https://github.com/marbl/CHM13. The data for HG01243 genome were released by Shafin et al., and are available from https://github.com/human-pangenomics/hpgp-data.

## References

1. Bashir A, Klammer AA, Robins WP, Chin CS, Webster D, Paxinos E, Hsu D, Ashby M, Wang S, Peluso P, Sebra R. A hybrid approach for the automated finishing of bacterial genomes. Nature biotechnology. 2012 Jul;30(7):701–7.

2. Boetzer M, Pirovano W. SSPACE-LongRead: scaffolding bacterial draft genomes using long read sequence information. BMC bioinformatics. 2014 Dec;15(1):1–9.

3. Warren RL, Yang C, Vandervalk BP, Behsaz B, Lagman A, Jones SJ, Birol I. LINKS: Scalable, alignment-free scaffolding of draft genomes with long reads. GigaScience. 2015 Dec 1;4(1):s13742–015.

4. Qin M, Wu S, Li A, Zhao F, Feng H, Ding L, Ruan J. LRScaf: improving draft genomes using long noisy reads. BMC genomics. 2019 Dec;20(1):1–2.

5. Li H. Minimap2: pairwise alignment for nucleotide sequences. Bioinformatics. 2018 Sep 15;34(18):3094–100.

6. Kolmogorov M, Yuan J, Lin Y, Pevzner PA. Assembly of long, error-prone reads using repeat graphs. Nature biotechnology. 2019 May;37(5):540–6.

7. Lawler EL. Combinatorial optimization: networks and matroids. Courier Corporation; 2001.

8. Lee H, Gurtowski J, Yoo S, Marcus S, McCombie WR, Schatz M. Error correction and assembly complexity of single molecule sequencing reads. BioRxiv. 2014 Jan 1:006395.

9. Berlin K, Koren S, Chin CS, Drake JP, Landolin JM, Phillippy AM. Assembling large genomes with single-molecule sequencing and locality-sensitive hashing. Nature biotechnology. 2015 Jun;33(6):623.

10. Gurevich A, Saveliev V, Vyahhi N, Tesler G. QUAST: quality assessment tool for genome assemblies. Bioinformatics. 2013 Apr 15;29(8):1072–5.

11. Miga KH, Koren S, Rhie A, Vollger MR, Gershman A, Bzikadze A, Brooks S, Howe E, Porubsky D, Logsdon GA, Schneider VA. Telomere-to-telomere assembly of a complete human X chromosome. Nature. 2020 Sep;585(7823):79–84.

12. Nurk S, Koren S, Rhie A, Rautiainen M, Bzikadze AV, Mikheenko A, Vollger MR, Altemose N, Uralsky L, Gershman A, Aganezov S. The complete sequence of a human genome. bioRxiv. 2021 Jan 1.

13. Zimin AV, Marçais G, Puiu D, Roberts M, Salzberg SL, Yorke JA. The MaSuRCA genome assembler. Bioinformatics. 2013 Nov 1;29(21):2669–77.

14. Zimin AV, Puiu D, Luo MC, Zhu T, Koren S, Marçais G, Yorke JA, Dvořák J, Salzberg SL. Hybrid assembly of the large and highly repetitive genome of Aegilops tauschii, a progenitor of bread wheat, with the MaSuRCA mega-reads algorithm. Genome research. 2017 May 1;27(5):787–92.

15. Shafin K, Pesout T, Lorig-Roach R, Haukness M, Olsen HE, Bosworth C, Armstrong J, Tigyi K, Maurer N, Koren S, Sedlazeck FJ. Nanopore sequencing and the Shasta toolkit enable efficient de novo assembly of eleven human genomes. Nature biotechnology. 2020 Sep;38(9):1044–53.

